# Adipocyte JAK2 Mediates Growth Hormone-Induced Hepatic Insulin Resistance

**DOI:** 10.1101/076265

**Authors:** Kevin C. Corbit, João Paulo G. Camporez, Jennifer L. Tran, Camella G. Wilson, Dylan Lowe, Sarah Nordstrom, Kirthana Ganeshan, Rachel J. Perry, Gerald I. Shulman, Michael J. Jurczak, Ethan J. Weiss

**Affiliations:** Cardiovascular Research Institute, University of California, San Francisco, San Francisco, CA USA, 94143; Department of Internal Medicine, New Haven, CT USA, 06520; Department of Cellular and Molecular Physiology, New Haven, CT USA, 06520; Howard Hughes Medical Institute, Yale University School of Medicine, New Haven, CT USA, 06520; Department of Medicine, Division of Endocrinology and Metabolism, University of Pittsburgh, Pittsburgh, PA USA, 15261

## Abstract

For nearly 100 years, Growth Hormone (GH) has been known to impact insulin sensitivity and risk of diabetes. However, the tissue governing the effects of GH signaling on insulin and glucose homeostasis remains unknown. Excess GH reduces fat mass and insulin sensitivity. Conversely, GH insensitivity (GHI) is associated with increased adiposity, augmented insulin sensitivity, and protection from diabetes. Here we induce adipocyte-specific GHI through conditional deletion of *Jak2* (JAK2A), an obligate transducer of GH signaling. Similar to whole-body GHI, JAK2A mice had increased adiposity and extreme insulin sensitivity. Loss of adipocyte *Jak2* augmented hepatic insulin sensitivity and conferred resistance to diet-induced metabolic stress without overt changes in circulating fatty acids. While GH injections induced hepatic insulin resistance in control mice, the diabetogenic action was absent in JAK2A mice. Adipocyte GH signaling directly impinged on both adipose and hepatic insulin signal transduction. Collectively, our results show that adipose tissue governs the effects of GH on insulin and glucose homeostasis. Further, we show that JAK2 mediates liver insulin sensitivity via an extra-hepatic, adipose tissue-dependent mechanism.

## INTRODUCTION

Argentinian Physician Scientist, Bernardo Houssay demonstrated that injection of anterior pituitary extract worsens glycemic control in dogs (1, 2). In contrast, loss of anterior pituitary function leads to hypoglycemia and increased sensitivity to insulin (3). Similar results were observed in humans where hypophysectomy ameliorates not only insulin resistance (4–6) but diabetic complications as well (7, 8). More recently, it has been demonstrated that Growth Hormone (GH) is responsible for much of the pituitary-derived diabetogenic activity (9). Both loss- and gain-of-function studies in humans and rodents support a role for GH in the biology of insulin responsiveness. Specifically, loss of GH receptor (GHR) function in humans and mice is associated with insulin sensitivity and protection against age-related diabetes (10, 11). Conversely, acromegalic patients with excessive GH secretion and transgenic *Gh* over-expressing mice have increased mortality and insulin resistance (12–14). Collectively, robust physiologic and genetic data support a prominent role for GH signaling in insulin/glucose homeostasis and the etiology of diabetes.

One of the major physiological functions of GH is controlling adipose tissue lipolysis (15, 16). Recent studies have demonstrated a critical role for insulin-mediated suppression of adipocyte lipolysis in the acute inhibition of hepatic gluconeogenesis through reductions in hepatic acetyl-CoA, leading to decreased pyruvate carboxylase activity and flux. Further, macrophage-induced lipolysis was shown to promote increased rates of hepatic gluconeogenesis and fasting hyperglycemia by promoting increased hepatic acetyl CoA content and pyruvate carboxylase activity/flux, as well as increased conversion of glycerol to glucose (17). Although the ability for GH to promote lipolysis directly is ambiguous, data supporting a causal role for lipolysis in GH-mediated insulin resistance are robust (18, 19). The role, if any, of lipolytic activity in Growth Hormone Deficiency or GHI-associated augmentation of insulin sensitivity is entirely unknown.

The tissue(s) mediating the effects of GH signaling on insulin and glucose homeostasis has been elusive. Loss of GH signaling in liver through conditional deletion of the GH receptor *Ghr* (20), *Stat5 (21)*, or *Jak2* (22) confers lean body mass, fatty liver, and insulin resistance. In stark contrast, mice (23) and humans (10) with global disruption of GHR have increased adiposity and insulin sensitivity.

There are conflicting results regarding skeletal muscle GHI on whole-body insulin sensitivity (24, 25), none of which phenocopy global GHI. Mice with beta-cell specific disruption of GHR show little effect on fasting insulin levels or insulin content in the pancreas on normal chow (26). Recently, mice with fat-specific disruption of GHR were described to have increased fat mass but no change in insulin or glucose homeostasis (27). However, these mice were generated using the *Fabp4:Cre* which is known to have activity outside of adipose tissue (28).

In an effort to determine the specific role of adipose tissue in the metabolic activity of GH, we deleted *Jak2* from adipocytes using *Adiponectin:Cre* (here termed JAK2A) (29). Similar to whole-body GH insensitivity (GHI), and as described earlier the resulting JAK2A mice had increased adiposity. Yet they also had improved whole-body insulin sensitivity. Interestingly, while chronic systemic GH exposure promoted hepatic insulin resistance and lipolysis in control mice, JAK2A animals were refractory to the diabetogenic action of GH. Prominent mechanisms regulating hepatic insulin sensitivity, including reductions in free fatty acids and liver lipid content, failed to account for the insulin sensitizing effects observed in JAK2A mice. Collectively, our work demonstrates that adipose tissue regulates the diabetogenic activity of GH and that a JAK2- dependent, adipose-derived factor mediates whole-body insulin sensitivity.

## RESULTS

**Adipocyte-specific deletion of *Jak2* augments adiposity and insulin responsiveness.** We disrupted *Jak2* specifically in adipocytes using *Adiponectin:Cre* on an inbred C57BL/6 background (30). All studies were done using male mice. On normal chow, body weight between control and JAK2A cohorts did not differ (Figure 1A). However, JAK2A mice had increases in both absolute and percent body fat (Figures 1B and 1C). Both visceral epididymal and subcutaneous inguinal fat pad mass were higher in JAK2A animals (Figures 1D and 1E), which was accompanied by adipocyte hypertrophy (Supplemental Figures 1A and 1B) without changes in immunological populations (Supplemental Figures 1C-1F, Supplemental Table 1) or circulating IL-6 (Supplemental Figure 1G. Despite increased adiposity, fasting JAK2A mice were hypoglycemic and trended toward hypoinsulinemia (Figures 1F and 1G), implying increased whole-body insulin sensitivity. Consistent with this, the JAK2A cohort was hyper-responsive to insulin during the insulin tolerance test (ITT, Figure 1H). Thus, despite increased adiposity JAK2A mice have augmented whole-body insulin responsiveness.

**FIGURE 1.**
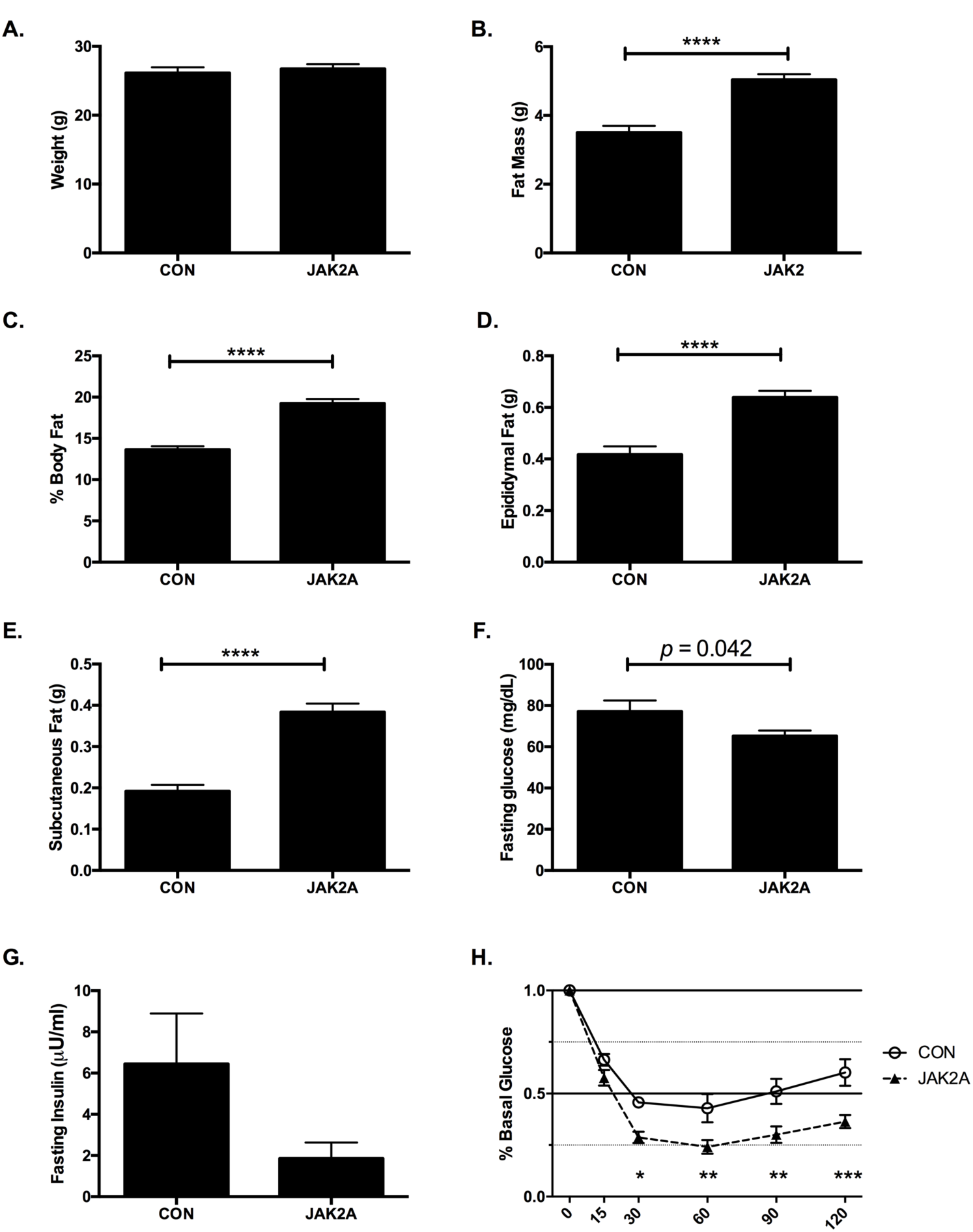
Loss of adipocyte *Jak2* augments insulin responsiveness in chow fed mice. a.) Total body weight and b.) fat mass in control (CON) and JAK2A mice. c.) Percent body fat as a fraction of total body weight. d.) Total epididymal visceral and e.) inguinal subcutaneous fat. f.) Fasting plasma glucose and g.) insulin. h.) Insulin tolerance testing expressed as a percent of basal (fasting) glucose in control (open circles) and JAK2A (black triangles) mice. *p<0.05, **p<0.01, ***p<0.001, ****p<0.0001 by T test (b-f) and OneWay ANOVA (h). (g)=grams. N=7-11 for both cohorts. Data are +/- S.E.M.

**Increased hepatic insulin sensitivity and suppression of endogenous glucose production in JAK2A mice.** To definitively determine tissue-specific insulin sensitivity, hyperinsulinemic-euglycemic clamps were performed. There was no difference in clamped plasma insulin levels between groups during the clamp (Supplemental Figure 2B) and plasma glucose levels were matched at approximately 120 mg/dl (Figure 2A). As compared to the control cohort, JAK2A animals required a higher glucose infusion rate (GIR) to maintain euglycemia, confirming augmented whole-body insulin sensitivity (Figures 2B and 2C). While basal endogenous glucose production (EGP) was unchanged between groups (Figure 2D), clamped JAK2A mice suppressed EGP following insulin infusion nearly 100% and to a much greater extent than control animals (Figures 2E and 2F), demonstrating improved hepatic insulin sensitivity. Whole-body glucose disposal (Figure 2G) and skeletal muscle- (Figure 2H) and adipose tissue-specific (Figure 2I) glucose uptake did not differ between the cohorts. Therefore, disruption of adipocyte *Jak2* confers improved whole-body insulin sensitivity almost entirely via augmented hepatic insulin sensitivity and suppression of hepatic glucose production.

**FIGURE 2.**
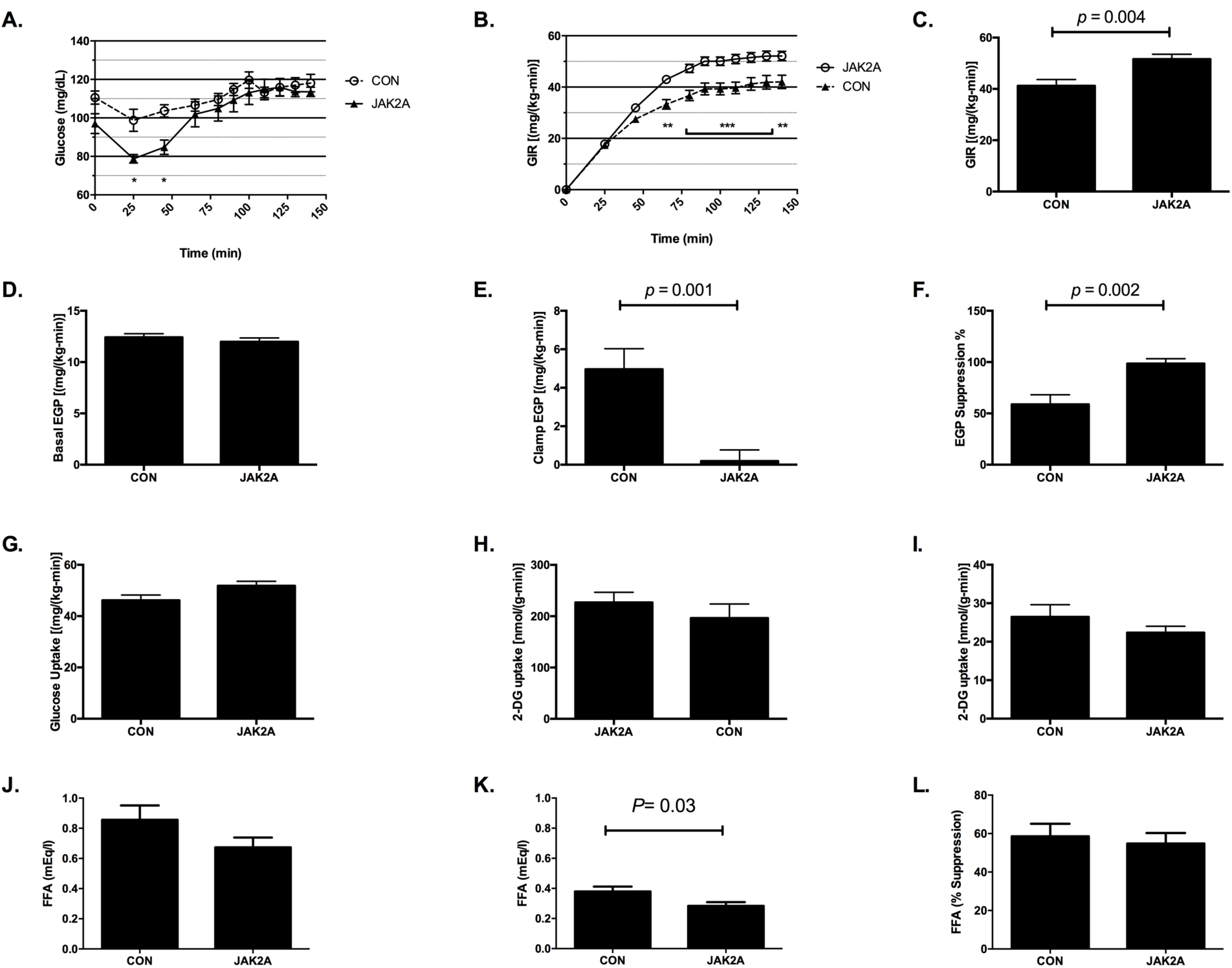
Adipocyte *Jak2* mediates hepatic insulin sensitivity in chow fed mice. a.) Blood glucose levels and b.) glucose infusion rate (GIR) during glucose infusion to achieve euglycemia in control (CON, open circles) and JAK2A (black triangles) mice. c.) GIR at euglycemia in control and JAK2A mice. d.) Basal and e.) clamped endogenous glucose production (EGP) in control and JAK2A mice. f.) Percent suppression of EGP following insulin infusion in control and JAK2A mice. g.) Whole-body glucose uptake in control and JAK2A mice. h.) 2-deoxyglucose (2-DG) uptake in gastrocnemius and i.) epididymal visceral fat in control and JAK2A mice. j.) Basal and k.) clamped plasma free fatty acid (FFA) levels in control and JAK2A mice. l.) Percent suppression of plasma FFA following insulin infusion in control and JAK2A mice. *p<0.05, **p<0.01, ***p<0.001 by OneWay ANOVA (a and b) and T test (c, e, f, and k). N=9 for both cohorts. Data are +/- S.E.M.

**Enhanced insulin-mediated suppression of lipolysis in JAK2A mice.** It was recently shown that reductions in adipocyte lipolysis and fatty acids (FA) are major regulators of insulin-mediated suppression of hepatic glucose output (31, 32). Since GH is a major in vivo regulator of lipolysis and loss of adipocyte *Jak2* greatly augmented insulin-induced reductions in EGP, we examined FA levels in the clamped mice. Plasma FA did not statistically differ between the euglycemic control and JAK2A cohorts (Figure 2J). Following insulin infusion the JAK2A cohort had lower absolute plasma FA (Figure 2K). In addition, insulin-mediated attenuation of isoproterenol-stimulated lipolysis was augmented in JAK2A adipose explants (Supplemental Figure 2H). However, no difference was appreciated between control and JAK2A mice when expressed as a percent suppression of basal FA (Figure 2L). Thus, while reduced lipolysis may contribute to augmented hepatic insulin sensitivity in JAK2A mice, the magnitude of difference between insulin-mediated suppression of EGP and FA (compare Figures 2E and 2K) presents the possibility of potential alternative mechanisms.

**JAK2A mice are resistant to diet-induced metabolic derangement.** To determine the susceptibility to metabolic stress, we next challenged mice with a high-fat diet (HFD). After ten weeks of HFD, body weight (Figure 3A), total (Figure 3B) and percent (Figure 3C) fat mass, and epididymal fat mass (Figure 3D) were unchanged between the control and JAK2A cohorts. The inguinal subcutaneous fat pads weighed more in JAK2A mice (Figure 3E). Both fasting blood glucose and insulin were reduced in JAK2A animals (Figures 3F and 3G). During ITT the JAK2A cohort maintained remarkable insulin responsiveness, suggesting whole-body insulin sensitivity was preserved despite HFD (Fig 3H).

**FIGURE 3.**
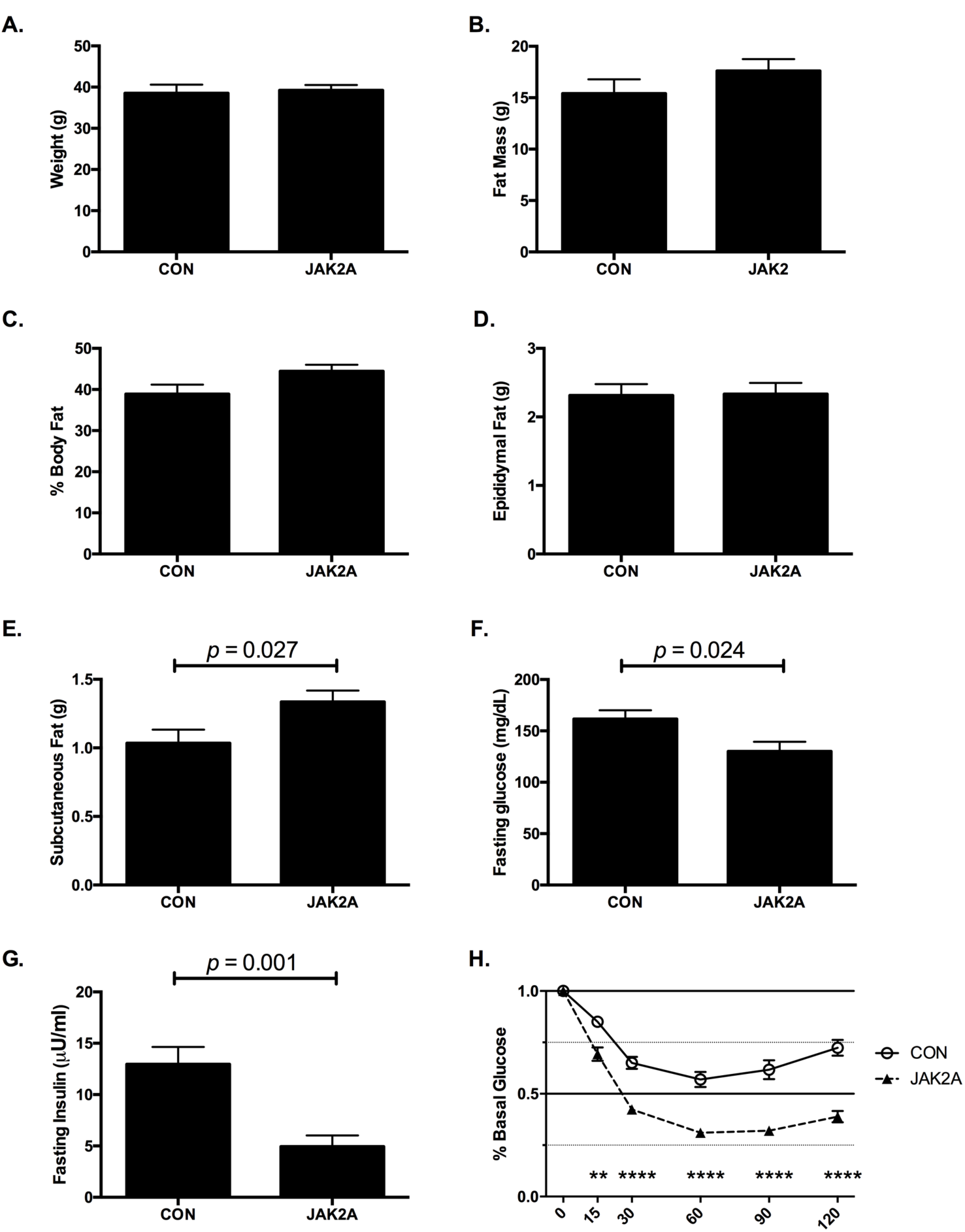
JAK2A mice maintain exquisite insulin responsiveness following dietary challenge. a.) Total body weight and b.) fat mass in control (CON) and JAK2A mice. c.) Percent body fat as a fraction of total body weight. d.) Total epididymal visceral and e.) inguinal subcutaneous fat. f.) Fasting plasma glucose and g.) insulin. h.) Insulin tolerance testing expressed as a percent of basal (fasting) glucose in control (open circles) and JAK2A (black triangles) mice. **p<0.01, **** p<0.0001 by T test (e, f, and g) and OneWay ANOVA (h). (g)=grams. N=8-16 for both cohorts. Data are +/- S.E.M.

**JAK2A mice maintain exquisite hepatic insulin sensitivity following HFD without changes in FA levels.** Hyperinsulinemic-euglycemic clamp experiments showed conclusively that JAK2A mice retained exquisite insulin sensitivity following HFD. Plasma insulin (Supplemental Figure 2D) and glucose levels (Fig. 4A) were matched in response to a fixed and variable rate infusion, respectively, during the clamp. At steady-state, GIR was nearly doubled in JAK2A animals (Figure 4C). Similar to mice on chow, basal EGP was unchanged between the groups (Figure 4D). However, while clamped control mice demonstrated hepatic insulin resistance, EGP was completely suppressed following insulin infusion in JAK2A animals, and furthermore appeared negative (Figures 4E and 4F). Negative EGP as measured by tracer dilution during a hyperinsulinemic euglycemic clamp is a well-documented phenomenon that typically occurs in extremely insulin sensitive models with high rates of glucose turnover, likely due to variable mixing of the exogenous 3-^3^H-glucose tracer with the endogenous glucose pool (33). Variable mixing of the 3-^3^H-glucose tracer also impacts measures of whole-body glucose disposal, which appeared unchanged between the two cohorts (Figure 4G). However, plasma clearance and tissue accumulation of the non-metabolizable glucose tracer ^14^C-2-deoxyglucose (2-DG) is not impacted by variable mixing and is therefore a more direct and reliable measure of glucose disposal. Tissue-specific glucose transport determined by 2-DG clearance was significantly increased in both skeletal muscle (Figure 4H) and adipose tissue (Figure 4I) in JAK2A mice. These data demonstrate that, despite the technical limitations of the 3-^3^H-glucose tracer under the conditions used, both hepatic and peripheral insulin sensitivity were improved in HFD JAK2A mice. In contrast, basal (Figure 4J) and insulin-mediated suppression of plasma FA (Figures 4K and 4L) did not differ between control and JAK2A animals. Therefore, adipocyte *Jak2* mediates whole-body insulin sensitivity even when challenged with HFD. Further, JAK2A mice augmented hepatic insulin sensitivity independent of statistically significant changes in FA levels. Collectively, JAK2A mice are resistant to the metabolic derangements of HFD.

**FIGURE 4.**
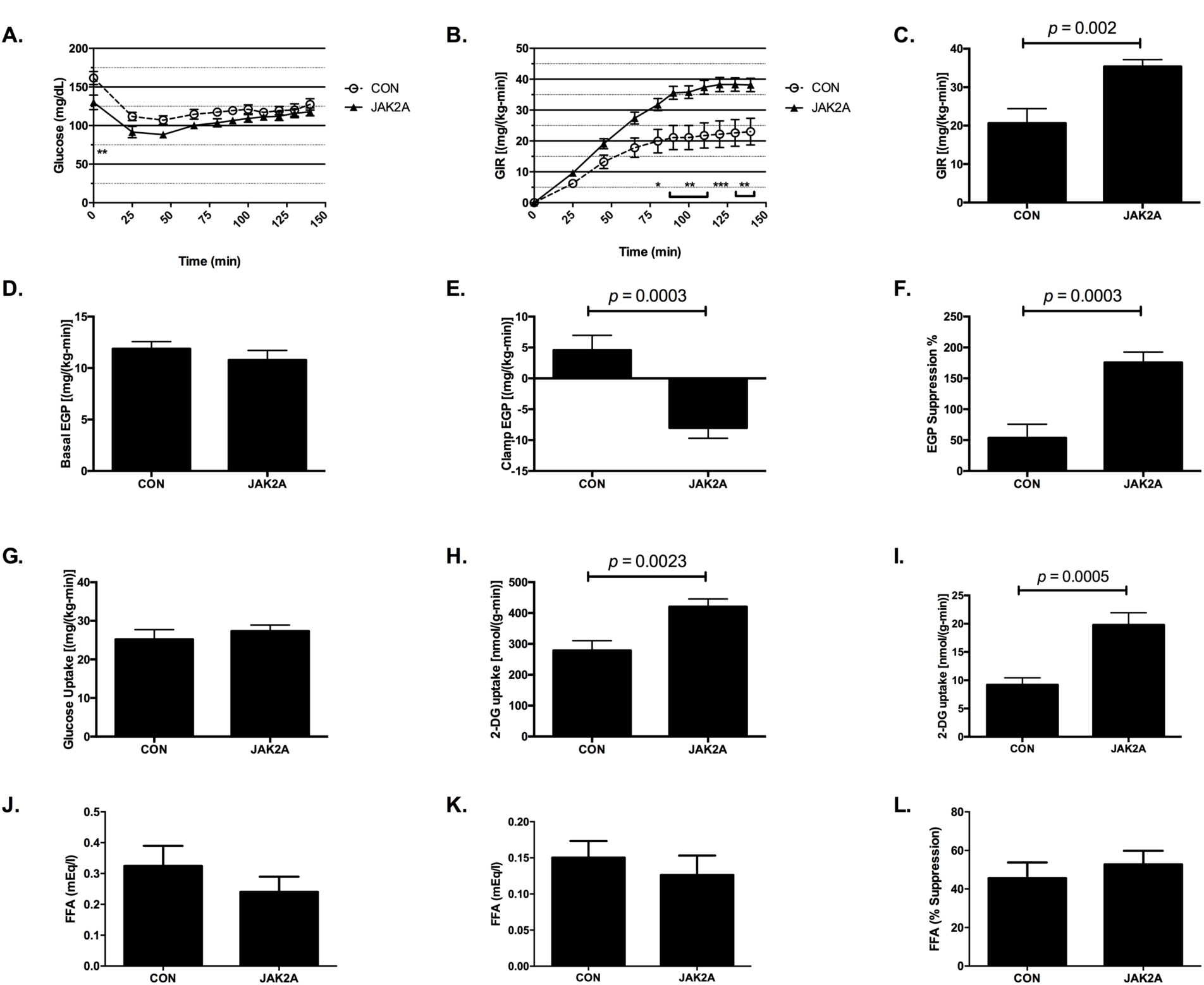
Loss of adipocyte *Jak2* imparts hepatic and whole-body insulin sensitivity in high fat diet fed mice. a.) Plasma glucose levels and b.) glucose infusion rate (GIR) during glucose infusion to achieve euglycemia in control (CON, open circles) and JAK2A (black triangles) mice. c.) GIR at euglycemia in control and JAK2A mice. d.) Basal and e.) clamped endogenous glucose production (EGP) in control and JAK2A mice. f.) Percent suppression of EGP following insulin infusion in control and JAK2A mice. g.) Whole-body glucose uptake in control and JAK2A mice. h.) 2-deoxyglucose (2-DG) uptake in gastrocnemius and i.) epididymal visceral fat in control and JAK2A mice. j.) Basal and k.) clamped plasma fatty acid (FA) levels in control and JAK2A mice. l.) Percent suppression of plasma FA following insulin infusion in control and JAK2A mice. *p<0.05, **p<0.01, ***p<0.001 by OneWay ANOVA (a and b) and T test (c, e, f, h, and i). N=8-11 for both cohorts. Data are +/- S.E.M.

**Reduced hepatic triacylglycerol levels do not account for increased insulin sensitivity in chow fed JAK2A mice.** Increased tissue lipid deposition strongly correlates with insulin resistance. Therefore, we reasoned that livers of JAK2A mice would have reduced lipid burden. Surprisingly, total levels of hepatic triacylglycerol (TAG) were unaltered between control and JAK2A chow fed cohorts (Supplemental Figure 2E). Therefore hepatic insulin sensitivity in chow-fed JAK2A mice is not associated with reduced total liver lipid content. Following HFD hepatic TAG content was increased in both control and JAK2A animals (Supplemental Figure 2E). Hepatic TAG levels in HFD-fed JAK2A mice were lower than the control cohort and hence may contribute to preserved insulin sensitivity in JAK2A animals following dietary challenge.

**Acute GH treatment induces hepatic insulin resistance which is dependent on adipocyte *Jak2***. Having established that insulin sensitivity is augmented in the setting of loss-of-function JAK2A mutants, we next determined the role of adipose tissue *Jak2* in gain-of-function GH signaling. To this end, control and JAK2A mice were treated with vehicle or supra-physiological doses of recombinant mouse GH daily for five days. GH treatment induced hyperinsulinemia in control and, to a lesser extent, in JAK2A mice (Supplemental Figure 2F). Following the hyperinsulinemic clamp, plasma insulin levels were equivalent in all groups (Supplemental Figure 2G). Similar to our previous chow-fed clamp study (Figure 2), vehicle-injected JAK2A mice required a higher GIR compared to the control cohort (Figures 5B and 5C). Systemic GH treatment reduced GIR by ~35% in control animals and by ~19% in JAK2A mice (Figure 5C). Basal EGP did not differ between groups (Figure 5D). Consistent with our previous study, JAK2A vehicle-treated mice demonstrated augmented EGP suppression following insulin infusion (Figure 5F). Conversely, GH treatment significantly reduced insulin-mediated EGP suppression in control animals, confirming the diabetogenic activity of GH (Figure 5F). Remarkably, GH injections had no effect on hepatic insulin sensitivity in the JAK2A cohort (Figures 5E and 5F). GH treatment diminished whole-body glucose disposal (Figure 5G) and skeletal muscle glucose uptake (Figure 5H) in both control and JAK2A mice. No differences in adipose tissue glucose uptake were observed (Figure 5I). Thus, only five days of systemic GH exposure induces hepatic insulin resistance in an adipocyte Jak2-dependent manner.

**FIGURE 5.**
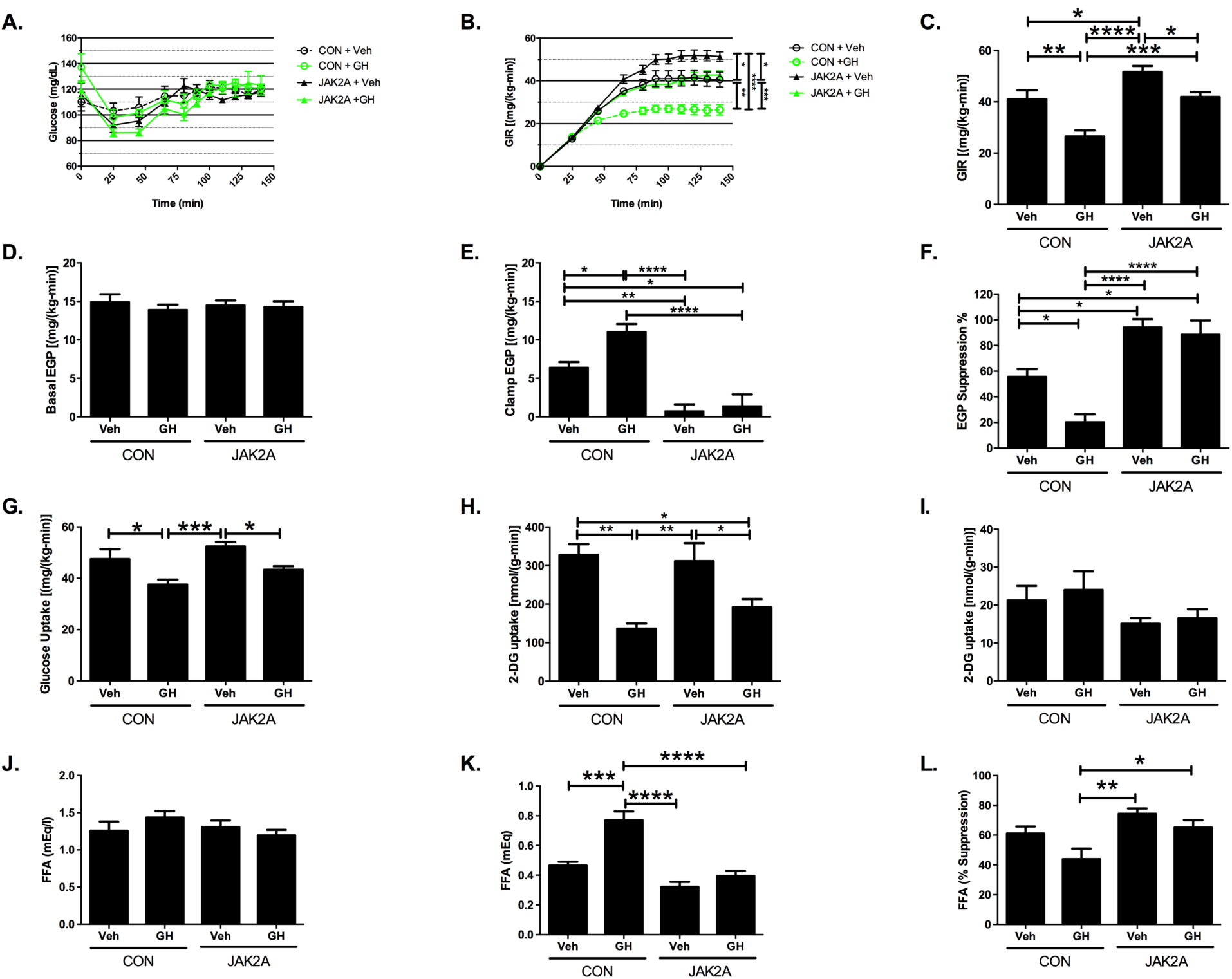
Adipocyte *Jak2* is required for Growth Hormone-induced hepatic insulin resistance and lipolysis. a.) Plasma glucose levels and b.) glucose infusion rate (GIR) during glucose infusion to achieve euglycemia in vehicle (Veh)-injected control (CON, open black circles) and JAK2A (closed black triangles) mice and Growth Hormone (GH)-injected control (open green circles) and JAK2A (closed green triangles) cohorts. c.) GIR at euglycemia in vehicle and GH-injected control and JAK2A mice. d.) Basal and e.) clamped endogenous glucose production (EGP) in vehicle- and GH-injected control and JAK2A mice. f.) Percent suppression of EGP following insulin infusion in vehicle- and GH-injected control and JAK2A mice. g.) Whole-body glucose uptake in vehicle- and GH-injected control and JAK2A mice. h.) 2-deoxyglucose (2-DG) uptake in gastrocnemius and i.) epididymal visceral fat in vehicle- and GH-injected control and JAK2A mice. j.) Basal and k.) clamped plasma free fatty acid (FFA) levels in control and JAK2A mice. l.) Percent suppression of plasma FFA following insulin infusion in control and JAK2A mice. *p<0.05, **p<0.01, ***p<0.001, ****p<0.0001 by OneWay ANOVA (a and b). N=6-8 for both cohorts. Data are +/- S.E.M.

**GH promotes adipocyte lipolysis indirectly via JAK2-dependent inhibition of insulin action.** In hyperinsulinemic-euglycemic clamped mice, circulating basal FA levels were unchanged in GH-injected animals (Figure 5J). Following insulin infusion, GH-injected control animals failed to suppress plasma FA to the levels of vehicle-treated mice, revealing that GH promoted lipolysis indirectly via inhibitory effects on insulin activity (Figure 5K). The ability of GH to antagonize reductions in circulating FA was absent in JAK2A mice (Figures 5K and 5L), demonstrating that adipocyte *Jak2* transduces the GH signal to interfere with insulin-mediated suppression of lipolysis. This correlated with retention of hepatic insulin sensitivity (Figure 5E). Collectively, we conclude that adipose tissue mediates GH-dependent antagonism of insulin’s activity in both hepatocytes and adipocytes.

Similar to our earlier clamp experiment (Figure 2), neither basal (Figure 5J) nor percent suppression of circulating FFA (Figure 5L) differed between vehicle treated control and JAK2A animals. Again, the lack of a demonstrable effect on lipolysis occurred concurrent with augmented hepatic insulin sensitivity (Figure 5E). A nearly identical pattern was observed for palmitate turnover (Supplemental Figures 2I-2K). Therefore, hepatic insulin resistance in GH-treated control animals directly correlates with adipocyte Jak2-dependent lipolysis. Further, the augmented hepatic insulin sensitivity of JAK2A mice may be mediated by a mechanism(s) other than or in addition to inhibition of lipolysis.

**GH treatment impairs adipocyte insulin responsiveness.** Since GH attenuated insulin-mediated suppression of lipolysis (Figure 5K), we determined the ability of insulin to induce signal transduction in adipose tissue from GH treated mice. Intraperitoneal injection of insulin induced robust insulin receptor (IR) autophosphorylation in control inguinal adipose tissue following a ten-minute exposure (Supplemental Figures 3A and 3B). Daily GH injections for seven days potently attenuated IR autophosphorylation in control but not JAK2A mice. Similarly, GH inhibited insulin induced AKT phosphorylation (pAKT) in a JAK2-dependent manner (Supplemental Figures 3C and 3D). Finally, and consistent with a previous report (34), *Pik3r1* (p85α) levels were reduced in JAK2A inguinal adipose tissue.

**Adipocyte *Jak2* governs hepatic insulin responsiveness.** Given that neither liver lipid content nor lipolysis strongly correlated with hepatic insulin sensitivity in mice lacking adipocyte *Jak2,* we measured hepatic insulin signaling in control and JAK2A cohorts. To this end, we continually exposed mice to recombinant GH for 28 days via mini-osmotic pumps. Subsequently, hepatic insulin responsiveness was assessed via levels of pAKT following injections into the inferior vena cava (i.v.c). Vehicle infused control animals responded robustly to i.v.c. insulin (Figure 6). Conversely, chronic GH exposure strongly abrogated insulin-induced hepatic pAKT, consistent with the diabetogenic action of GH observed during the clamp studies (Figure 5) and previous work (35). However, GH treatment did not impair insulin-induced pAKT in JAK2A livers, demonstrating that adipocyte JAK2 mediates GH-induced decrease in hepatic insulin responsiveness.

**FIGURE 6.**
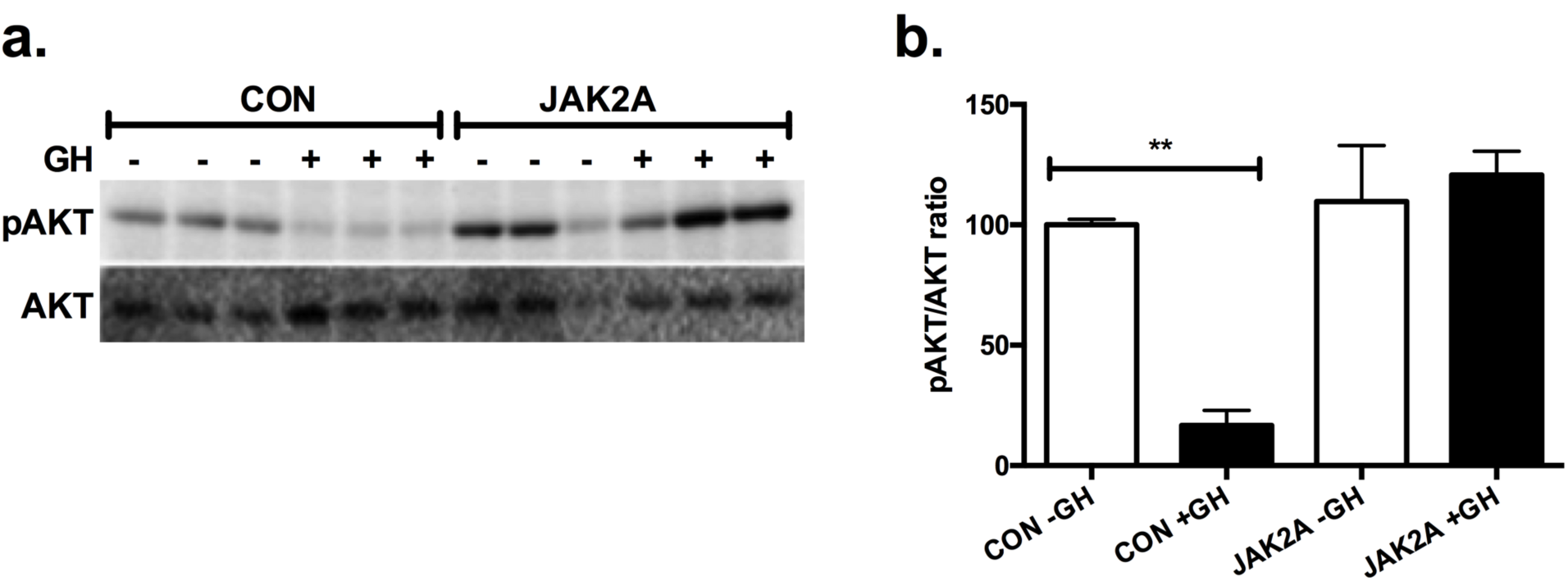
Adipocyte *Jak2* mediates insulin-induced hepatic AKT activation. a.) Western blot of insulin-stimulated liver lysates with antibodies against phosphorylated (pAKT) and total AKT (AKT) in control and JAK2A mice following continuous delivery of vehicle (-) or recombinant Growth Hormone (GH, +) for 28 days. d.) Densitometric ratio of pAKT/AKT in control and JAK2A mice treated (+GH) or not (-GH) with GH. *p<0.05, **p<0.01, ****p<0.0001 by OneWay ANOVA. Data are +/- S.E.M.

## DISCUSSION

Houssay and colleagues first described their landmark studies on extreme insulin sensitivity in hypophysectomized dogs nearly 100 years ago (1). In 1987, Mayer Davidson lamented that "the mechanism of GH effects on carbohydrate and lipid metabolism is far from clear.” Further, "Several studies suggest that the insulin antagonistic effects of GH may be secondary to the increased lipolysis via the glucose-fatty acid cycle. It is disappointing to note that this was the conclusion of the last general review of this area more than 20 years ago, and we have made little progress in substantiating or refuting this hypothesis (36)." Now, almost 30 years after this review, there is still no clear understanding of the cellular or tissue-level basis underlying the effects of GH on insulin and glucose homeostasis.

Global GHI, in both the physiologic and genetic context, confers insulin sensitivity throughout life. Conversely, acromegalic patients with excessive GH secretion are prone to insulin resistance, risk of diabetes and increased mortality. In fact, it has been proposed that GH hypersecretion may be the cause as much as the consequence of poor diabetic control (37). These effects have been well documented in dogs, mice, and teleosts (38), demonstrating robust evolutionary conservation. However, the vast array of loss-of-function studies describing inactivation of GHR, JAK2, or STAT5 in metabolic tissues including liver, skeletal muscle, pancreatic beta cell, and even adipose tissue has yielded complex and difficult to interpret results and failed to answer the key question of where GH signaling (at the cell or tissue level) mediates its effects on whole-body insulin and glucose metabolism. Here, we show conclusively that abrogation of GH signaling in adipocytes augments whole-body and hepatic insulin sensitivity, protects from HFD-induced whole-body and hepatic insulin resistance, and prevents GH-mediated hepatic insulin resistance.

Our work relies on JAK2 as a signaling intermediate to infer GH action. While there is clear evidence that GH signaling is entirely dependent on JAK2 in fat and other tissues, it is possible or even likely that JAK2 mediates signals downstream of other cytokine signaling receptors in fat. Conditional deletion of *Ghr* with AP2:Cre results in obesity with no improvements in glucose homeostasis (27). Removal of *Jak2* from adipose using the same *Cre* transgene was reported to also induce adiposity with age-related insulin resistance (39). This is in contrast to our JAK2A mice that, while obese, are exquisitely insulin sensitive, even when challenged with HFD or supra-physiologic amounts of GH. Our work here and previously (29) utilized *Adiponectin:Cre* which has been shown to be considerably more adipocyte-specific (40). The AP2:Cre line induces recombinase activity in the peripheral and central nervous systems as well as adrenal medulla (41). Therefore, loss of either *Ghr* or *Jak2* in these populations may mediate the published phenotypes using the AP2:Cre line. Thus, in the absence of an *Adiponectin:Cre-driven Ghr* deletion phenotype, it is not yet possible to conclude if the insulin sensitivity conferred by lack of adipose JAK2 is entirely due to interruption of GH signaling. However, we do definitively show that adipocyte JAK2 is required for GH to induce adipose tissue lipolysis and hepatic insulin resistance.

Insulin resistance strongly correlates with hepatic steatosis. Mechanistically, it has been shown that the subcellular localization of diacylglycerols (DAG) and not absolute levels of liver lipid mediate hepatic insulin resistance. Specifically, DAG and PKCε are membrane-associated in insulin resistant rats but lipid droplet bound in the insulin sensitive state, preventing PKCε translocation and activation at the plasma membrane where PKCε inhibits Insulin Receptor kinase activity through phosphorylation of threonine 1160 (42). These findings are germane, as in some cases liver lipid and hepatic insulin resistance do not correlate, such as CGI-58 ASO treatment (43). For instance, we recently published work demonstrating that resolution of hepatic steatosis in hepatocyte-specific *Jak2* knockout mice (JAK2L) via concomitant deletion of *Cd36* is not sufficient to reverse the insulin resistance observed in JAK2L animals (44). Others have shown discordance between insulin resistance and tissue lipid deposition, such as in Chanarin-Dorfman Syndrome and genetic forms of hepatic steatosis (45, 46).

Further, insulin resistance and adiposity fail to correlate in cases of Cowden and Laron Syndromes (47, 48). In fact, one of the natural consequences of augmented adipose tissue insulin sensitivity is reduced lipolysis, and hence increased adiposity, as is experienced by patients taking insulin-sensitizing agents such as thiazolidinedione (49). Here we report that chow-fed JAK2A have increased adiposity, but fail to demonstrate statistical differences in either basal or percent suppression of lipolysis following insulin infusion.

The physiological consequences of adipose tissue insulin sensitivity are glucose uptake and suppression of lipolysis. The former occurs via Akt-regulated GLUT4 transport (50). However, exactly how insulin suppresses adipose tissue lipolysis is unknown. The major facilitator of insulin-mediated inhibition of lipolysis appears to be PDE3B (51). Surprisingly, this has been reported to occur in an Akt-independent manner (52). While GH potently attenuated insulin-induced phosphorylation of Akt (Supplemental Figures 3C and 3D), no effects on adipose tissue glucose uptake were observed (Figure 5L). Therefore, GH seems to selectively affect the anti-lipolytic arm of adipose tissue insulin action. We were unable to detect adipose tissue PDE3B using commercially available antibodies, but a robust inquiry into the potential role of PDE3B in GH-mediated inhibition of insulin-induced suppression of lipolysis in warranted.

In 1963 Philip Randle put forth the notion that FAs may themselves induce insulin resistance (53). Indeed, increasing levels of circulating FAs highly correlate with insulin resistance (54). Further, clinical trials testing agents that reduce FAs have shown increased insulin sensitivity outcomes (55–58). One of the many actions of insulin is to inhibit adipose tissue lipolysis and hence FA release into the circulation. GH has also been proposed to be a major regulator of adipose tissue lipolysis (15). Indeed, acromegalic patients have enhanced lipolysis and increased adipose expression of genes that regulate lipolysis (59). Two independent trials reported that administration of the anti-lipolytic drug acipimox increases insulin sensitivity in GH-treated patients (18, 19). Collectively, these data support a predominant role for lipolysis in GH-mediated insulin resistance. Our work here supports these findings. We showed that GH-mediated lipolysis correlates with hepatic insulin resistance and that GH promoted adipose tissue lipolysis through JAK2, but indirectly via perturbation of insulin action. Therefore, the predominant diabetogenic mechanism of GH is likely due to increased adipose tissue lipolysis promoting hepatic insulin resistance.

However, it is unknown if abrogated adipose tissue lipolysis mediates the enhancement of insulin sensitivity observed under loss-of-function conditions, such as in Laron Syndrome and other forms of congenital GHI. As opposed to systemic GH treatment, our results do not support a role for modulation of lipolysis in JAK2A-associated insulin sensitivity. Specifically, we did not observe differences in circulating FA under clamped conditions between high fat diet fed control and JAK2A mice, despite augmented hepatic insulin sensitivity in the latter. This is consistent with clinical results reporting that inhibition of GH signaling in Type 1 diabetics improves hepatic insulin sensitivity without a clear effect on lipolysis (60). While the results from gain- and loss-of-function GH conditions appear disparate, the effects of GH on lipolysis may be determined by the nutritional state of the organism (61). Here, we present the possibility that the acromegalic and GHI states may mediate insulin sensitivity by different mechanisms. Alternatively, adipocyte JAK2 may antagonize insulin sensitivity by additional mechanisms independent of GH signaling.

Recently two groups reported that insulin-induced reductions in hepatic glucose output can occur independently of liver insulin signaling (31, 62). This surprising finding supports older literature showing that insulin inhibits hepatic glucose output indirectly via effects on adipose tissue (63–66) and suggests that extra-hepatic tissue(s) receive insulin signals to mediate hepatic insulin sensitivity. In support of this, selective enhancement of adipocyte insulin sensitivity imparts whole-body glucose homeostasis (67). Further, transplanting insulin sensitive adipose tissue into insulin resistance animals restores insulin responsiveness (68–70). Finally, overt lack of adipose tissue function is sufficient to induce insulin resistance (71). Therefore adipose tissue may be the dominant regulator of whole-body insulin sensitivity.

Exactly how adipose tissue mediates hepatic insulin sensitivity is unknown, but recent mechanisms have been proposed that include lipolysis (31, 62). We did not observe consistent differences in adipose tissue lipolysis or liver lipid content in chow-fed, HFD-fed or GH-treated insulin sensitized JAK2A mice. Further, we have not measured consistent or significant changes in classic insulin sensitizing factors such as Adiponectin and Leptin (29). Therefore, adipocyte JAK2 may mediate insulin sensitivity by a novel mechanism. Previously, it was reported the

GH mediates cellular insulin resistance at the level of PI 3-kinase by altering sub-cellular redistribution of AKT (72). We found that GH treatment attenuated insulin-induced pAKT in inguinal adipose tissue, but the molecular mechanism remains unknown.

GH has been reported to regulate p85α expression and PI 3-kinase activity in general (34). Consistently, we observed a reduction in *Pik3r1* levels in JAK2A adipose tissue (Supplemental Figure 3E), but are unable to comment on PI 3-kinase activity. When mutated in humans with Cowden Syndrome, the PI 3-kinase pathway effector *PTEN* promotes adiposity coupled to augmented insulin sensitivity (47).

Here, we report that loss of adipocyte *Jak2* is sufficient to impart whole-body insulin sensitivity independent of adiposity and liver lipid content, suggesting the presence of a paracrine regulator of GH diabetogenic activity and highlighting cross talk between adipose and liver. Our results show that acrogmegaly-associated hepatic insulin resistance is a result of attenuated insulin-mediated adipose tissue lipolysis. Finally, we provide evidence to support alternative mechanisms regulating gain- and loss-of-function GH signaling-mediated effects on insulin responsiveness. Isolation of the JAK2-dependent, adipocyte-derived factor(s) mediating whole-body insulin sensitivity may someday lead to new anti-diabetic therapeutics.

## METHODS

**Animals and diets.** The generation of *Adiponectin:CRE;Jak2^ox/lox^* mice was previously described (29). For our studies reported here, *Jak2^ox//ox^* (73))were used as controls and backcrossed onto the C57BL/6 background for at least nine generations. For chow fed studies, mice received PicoLab Mouse Diet 20 (Lab Diet #5058; percent calories provided by protein 23%, fat 22%, carbohydrate 55%) and for high fat diet studies mice were fed Research Diets D12492 (percent calories provided by protein 20%, fat 60%, carbohydrate 20%).

**Dietary studies.** Six week-old control and JAK2A mice were fed high fat diet or maintained on chow for ten weeks. Total fat mass was determined by Dual-energy X-ray absorptiometry. Blood glucose and serum insulin levels were determined by glucometer readings (Bayer Contour) and ELISA (Alpco), respectively, following an overnight (16 hours) fast. For insulin tolerance testing, mice were fasted for four hours (09:00-13:00) followed by intraperitoneal injection of 2 U/kg insulin (Novolin^®^ Novo Nordisk, Bagsvaerd Denmark). Blood glucose levels were determined by tail prick using a hand-held glucometer at the times indicated.

**Hyperinsulinemic-euglycemic clamp studies.** Clamp studies were performed according to recent recommendations of the NIH-funded Mouse Metabolic Phenotyping Consortium and as previously described (74, 75). Briefly, mice recovered one week after receiving surgery to implant an indewelling jugular vein catheter prior to clamp studies. Mice were fasted overnight and received a basal infusion of 3-^3^H-glucose and U-^13^C-palmitate conjugated to BSA to determine fasting glucose and palmitate turnover. A primed/continuous infusion of insulin, 3-^3^H-glucose and U-^13^C-palmitate was administered alongside a variable infusion of 20% dextrose to maintain euglycemia during the hyperinsulinemic portion of the study. A 10 µCi bolus injection of ^14^C-2-deoxyglucose was given at 90 min to determine tissue-specific glucose uptake, which was calculated from the area under the curve of ^14^C-2-deoxyglucose detected in plasma and the tissue content of ^14^C-2-deoxyglucose-6-phosphate. Blood was collected by tail massage at set intervals and glucose levels measured by a glucose oxidase method. Insulin infusion rates were as follows; chow 2.5 mUkg^-1^min^-1^; HFD 4.0 mUkg^-1^min^-1^; GH 2.5 mUkg^-1^min^-1^. Glucose and palmitate turnover were determined as the ratios of the 3-^3^H-glucose and U-^13^C-palmitate infusion rates to the specific activity or plasma enrichment corrected to the contribution of the infusion rate for plasma glucose and palmitate, respectively, during the last 40 min of the hyperinsulinemic infusion. Calculation of endogenouse glucose production and tissue specific glucose uptake were described previously (74). For chow studies mice were approximately 16 weeks of age and HFD study mice were age-matched to chow studies and fed HFD for 4 weeks prior to clamps. For GH treatment mice received daily vehicle (0.03M NaHCO3, 0.15 NaCl, pH 9.5) or 5 mg/kg recombinant mouse GH (Dr. A. F. Parlow, National Hormone and Peptide Program, UCLA, Torrance, CA) by subcutaneous injection for 5 days prior to clamp.

**Mini-osmotic pump studies.** Control and JAK2A mice were implanted with 28-day Alzet mini-osmotic pumps (Durect Corporation, Cupertino CA) to deliver recombinant mouse GH at a dose of 5 mg/kg/day in GH buffer. At the end of 28 days, mice were fasted overnight (16 hrs) followed by injection with 0.01U insulin via the inferior vena cava. After 5 minutes, the liver was removed and immediately snap frozen in liquid nitrogen. Liver tissue was prepared for gel electrophoresis as previously described (44). For detection of pAkt and total Akt, liver was homogenized in RIPA buffer (50 mM Tris, 150 mM NaCl, 1% Triton-X100, 1% deoxycholate, 0.1% SDS, 1mM EDTA, pH 7.4) supplemented with protease and phosphatase inhibitors (Thermo Fisher Scientific, Waltham MA). After incubating on ice for 20 minutes, the homogenate was centrifuged for 20 minutes at 15000 RCF at 4C. The protein concentration of the supernatant was determined using BCA assay (Pierce) and 20 µg of protein used for gel electrophoresis. Antibodies used were anti-phospho-S473-Akt (Cell Signaling Technologies #4060) and total Akt (Cell Si#9272) at a dilution of 1:2000 followed by anti-Rabbit secondary at a dilution of 1:10000. Images were collected using Chemidoc Imager (Bio-Rad, Hercules, CA).

**Statistics and graphics.** All statistical tests and figures were done using GraphPad Prism v6.0. For all T tests the P value is included in the figure and is considered significant if less than 0.05.

### Study approval

All animal studies were approved by IACUC at the Universities of California, San Francisco and Pittsburgh, and Yale University.

## Acknowledgements

This study was supported by National Institutes of Health (NIH) Grants 1R01DK091276 (to E.J.W.), DK076169 (M.J.J. & E.J.W.), DK099402 (M.J.J.), DK059635, DK40936, DK45735 (G.I.S.). We also gratefully acknowledge the support of the James Peter Read Foundation, the University of California, San Francisco (UCSF) Cardiovascular Research Institute, the UCSF Diabetes Center (P30 DK063720), and the UCSF Liver Center (P30 DK026743). We would like to thank Dr. Kay-Uwe Wagner from the University of Nebraska for kindly providing the *Jak2* conditional mice. We would also like to thank Dr. Ajay Chawla of UCSF for his advice and guidance.

## Author contributions

J. L. T. performed experiments for Figures 1 and 3 and the real-time qPCR. J. P. G. C. and R. J. P. executed experiments for Figures 2, 4, and 5. C. G. W. carried out experiments for Figure 6 and immune cell FACS analyses. D.L. did GH and insulin injections followed by adipose tissue Western blots. S. N. collected tissue and recorded microscopic pictures for adipose tissue histology, did the IL6 ELISAs and the adipose explant stimulated lipolysis assays. K. G. performed flow cytometric analysis of stromal adipose cells. G. I. S. provided critical insight and reviewed the manuscript. K. C. C. wrote the manuscript, performed statistical analyses, and made the figures. M. J. J. wrote the manuscript and designed and supervised the clamp experiments. E. J. W. conceived of the study, wrote the manuscript, performed statistical analyses, and made figures.

## Conflict of interest statement

The authors have declared that no conflict of interest exists.

